# Comparative Transcriptomic Analysis of a Kluyveromyces marxianus Chassis Expressing Heterologous Proteins with Different Expression Efficiencies

**DOI:** 10.64898/2026.11.26.769105

**Authors:** Wei Zhao, Haiyan Ren, Jungang Zhou, Yao Yu, Hong Lu

**Affiliations:** State Key Laboratory of Genetics and Development of Complex Phenotypes, School of Life Sciences, Fudan University, Shanghai 200438, China; Shanghai Engineering Research Center of Industrial Microorganisms, Shanghai 200438, China

## Abstract

*Kluyveromyces marxianus* (KM) is a promising host for recombinant protein production, yet host transcriptional responses associated with different protein expression efficiencies remain poorly understood. Here, we constructed an engineered KM chassis, FDHY232, by combining multiple expression-enhancing genetic traits and evaluated the production of five heterologous proteins. Human ferritin heavy chain achieved efficient intracellular soluble expression, reaching an estimated titer of 14.73 g/L in 5 L fed-batch cultivation. Feruloyl esterase and β-lactoglobulin variant B were efficiently secreted, whereas bovine lysozyme and human lactoferrin were poorly expressed and predominantly accumulated in intracellular insoluble fractions. Comparative transcriptomic analysis identified 18 candidate genes associated with protein expression efficiency and 29 candidate features distinguishing efficient intracellular expression from secretion. *ILV2* and *LYS21* were consistently expressed at higher levels in the efficiently expressed protein group, whereas *LAA1* and *MNN1* were associated with efficient secretion. Functional enrichment analysis revealed negative enrichment of translation and amino acid biosynthesis pathways in the poorly expressed protein group and positive enrichment of protein folding in the efficient secretion subgroup. These findings suggest that enhancing amino acid biosynthesis through *ILV2* and *LYS21*, maintaining translational capacity, and enhancing oxidative folding may improve the production of difficult-to-express proteins. This study provides insights into protein-dependent host transcriptional responses and potential engineering targets for rational chassis optimization in KM.

## Introduction

Recombinant proteins are widely used in biopharmaceuticals, industrial enzymes, and food biotechnology, and microbial cell factories have become an important platform for their production (1, 2). Compared with prokaryotic expression systems, yeasts combine genetic tractability and suitability for large-scale fermentation with the capacity for eukaryotic protein processing and secretion (1, 2). Among them, the non-conventional yeast Kluyveromyces marxianus (KM) has emerged as a promising host for heterologous protein production owing to its rapid growth, thermotolerance, and broad substrate utilization (3, 4). More than 50 recombinant proteins have been successfully expressed in KM (5), with production titers reaching 16.8 g/L (6). Recent advances in synthetic biology and genome engineering, including signaling pathway modulation, chromosomal rearrangement, and genome-wide rearrangement-based screening, have further improved recombinant protein titers in KM (7-9).

Despite these advances, heterologous proteins expressed in the same yeast chassis often exhibit substantial differences in expression efficiencies(10). While some proteins are efficiently produced intracellularly or secreted, others are poorly expressed or accumulate as insoluble aggregates. These differences may arise from both the intrinsic properties of target proteins, including amino acid sequence, molecular size, and folding and post-translational processing requirements, and host factors, such as cellular resource allocation, protein quality control, and secretory pathway capacity (11-13).

Transcriptomic analysis provides a powerful approach to understanding host responses to heterologous protein expression and identifying engineering targets. In *P. pastoris*, Gasser et al. identified and validated host helper factors that enhance recombinant protein secretion (14), while Baumann et al. identified protein trafficking and ergosterol metabolism as potential engineering targets (15). In *Y. lipolytica*, Korpys-Woźniak and Celińska revealed shared and protein-specific transcriptional responses associated with protein localization, cysteine content, and glycosylation characteristics (16). In KM, Shi et al. identified endoplasmic reticulum stress-responsive genes whose manipulation could modulate heterologous protein secretion (17), while Zhou et al. investigated the effects of *MNN11* deletion on protein secretion and host transcriptional responses, highlighting the role of glycosylation in recombinant protein production (18). However, previous transcriptomic studies in KM have primarily focused on specific host modifications and secretion-related processes. Systematic comparisons of host transcriptional responses to heterologous proteins with distinct production phenotypes within the same engineered KM chassis remain limited.

In this study, we constructed an engineered *K. marxianus* chassis, FDHY232, by combining multiple expression-enhancing genetic traits. Three heterologous proteins were efficiently expressed, with rhFTH reaching an estimated titer of 14.73 g/L, whereas two others remained poorly expressed. Comparative transcriptomic analysis identified candidate genes and functional pathways associated with distinct protein expression phenotypes, highlighting amino acid biosynthesis, translation, and protein folding as potential targets for further chassis optimization.

## 2. Materials and methods

### 2.1. Yeast strains

The *Kluyveromyces marxianus* strain T1-Δatg1, an *ATG1* deletion derivative of T1, was previously described by Liu et al. (19). An N1546K substitution was introduced into Cyr1 of T1-Δatg1 as previously described (7). Subsequently, Cre-loxP-mediated chromosomal translocation between loxP site 11 on chromosome III and loxP site 7 on chromosome V was performed as described by Ren et al. (8), yielding the engineered chassis strain FDHY232. Yeast strains were routinely cultivated at 30 °C in YPD medium (2% peptone, 1% yeast extract, and 2% glucose, supplemented with 2% agar for solid medium).

### 2.2. Amino acid composition analysis

Five heterologous proteins were selected for expression: human ferritin heavy chain 1 (rhFTH, UniProt ID: P02794), feruloyl esterase (AnFaeA, A2QSY5), β-lactoglobulin variant B (BLG-B, P02754), bovine lysozyme (bLYZ, Q06283), and human lactoferrin (hLF, P02788). To compare the amino acid composition of the five heterologous proteins with that of endogenous KM proteins, the KM reference proteome comprising 5,070 protein sequences was retrieved from UniProt. Sequence analysis and data visualization were performed using R (v4.5.2) in RStudio (v2026.01.0+392; Posit Software, PBC). The abundance and relative frequency of each of the 20 standard amino acids were calculated for each protein sequence. The distributions of amino acid frequencies across the KM reference proteome were visualized using boxplots, with the corresponding values of the five heterologous proteins superimposed for comparison.

### 2.3. Plasmids

pUKDN125-rhFTH (20),and pUKD-TEF-AnfaeA (LHZ766) (17) were used to express rhFTH and AnFaeA, respectively. The BLG-B coding sequence was amplified from LHZ1972 (21), and cloned between the SmaI and NotI sites of LHZ399 (22), to generate pUKDN132-C29T-BLG-B. For bLYZ and hLF, the native signal peptide-coding sequences were removed, and C-terminal His_6_ tags were introduced. The resulting coding sequences were codon-optimized, synthesized by GENEWIZ, and cloned into the same restriction sites of LHZ399, generating pUKDN132-C29T-bLYZ and pUKDN132-C29T-hLF, respectively. The codon-optimized sequences of bLYZ and hLF are provided in Supplementary File 1.

### 2.4. Heterologous protein expression in flasks

Expression plasmids were introduced into FDHY232 using the lithium acetate-mediated transformation method (23). Transformants were selected on synthetic complete medium lacking uracil (SC-Ura) (24). The recombinant strains expressing rhFTH, AnFaeA, BLG-B, bLYZ, and hLF were designated F-rhFTH, F-AnFaeA, F-BLG-B, F-bLYZ, and F-hLF, respectively. Five independent transformants were examined for rhFTH, AnFaeA, and BLG-B, whereas three independent transformants were examined for bLYZ and hLF. The recombinant strains were inoculated into 150-mL shake flasks containing 50 mL YG medium (2% yeast extract and 4% glucose) and cultivated at 30 °C and 220 rpm for 48 h.

For intracellular rhFTH analysis, cells from 1 mL of culture were harvested by centrifugation at 10,000 rpm for 5 min, washed with phosphate-buffered saline (PBS; 137 mM NaCl, 2.7 mM KCl, 10 mM Na_2_HPO_4_, and 1.8 mM KH_2_PO_4_, pH 7.4), resuspended in 500 μL PBS, and disrupted with 400 μL glass beads using a FastPrep-24 homogenizer (MP Biomedicals, Santa Ana, CA, USA). Cell lysates were centrifuged at 14,000 rpm for 20 min at 4 °C, and the supernatants were collected as intracellular soluble protein fractions for SDS-PAGE analysis.

For AnFaeA and BLG-B, culture supernatants were collected for SDS-PAGE analysis. For bLYZ and hLF, culture supernatants and intracellular soluble fractions were collected as described above. The insoluble pellets were thoroughly resuspended in SDS-PAGE loading buffer and boiled. After centrifugation, the resulting supernatants were collected as intracellular insoluble fractions.

For Western blot analysis of bLYZ and hLF, proteins were separated by SDS-PAGE, transferred onto PVDF membranes, and detected using a mouse anti-His antibody (1:5,000; M30111, Abmart, Shanghai, China) and an HRP-conjugated goat anti-mouse IgG secondary antibody (1:5,000; 074-1806, KPL, USA).

### 2.5. Fed-batch cultivation of rhFTH **and** F-BLG-B

Fed-batch cultivation was performed in 5 L bioreactors (BIOTECH-5BG-5-7500, BXBIO). Seed cultivation and feeding strategies were based on the method described by Chen et al. (25). Briefly, the temperature was maintained at 30 °C, and the pH was controlled at 5.5 by adding concentrated aqueous ammonia. Dissolved oxygen (DO) was regulated through cascade control of the agitation speed.

### 2.6. Semi-quantitative analysis of rhFTH and BLG-B titers

Protein titers during fed-batch cultivation were estimated by SDS-PAGE densitometry as described by Tian et al. (26) . Briefly, β-lactoglobulin standards (L3908, Sigma-Aldrich, USA) at concentrations of 0.1, 0.2, 0.3, 0.4, 0.5, and 1.0 g/L were separated alongside serially diluted protein samples on the same gel. After staining and destaining, integrated band intensities were quantified using ImageJ with local background subtraction. A calibration curve was generated by linear regression of β-lactoglobulin concentrations against background-corrected band intensities. rhFTH and BLG-B concentrations were estimated within the linear range of the calibration curve and corrected for dilution factors.

### 2.7. RNA extraction, library preparation, and sequencing

Cells were harvested after 48 h of shake-flask cultivation as described in Section 2.4. Samples comprised the parental strain FDHY232 and strains expressing the five heterologous proteins, with three independent biological replicates per group. Total RNA extraction, quality assessment using an Agilent 2100 Bioanalyzer, library preparation, and paired-end sequencing were performed by Novogene (China).

### 2.8. Bioinformatic analysis of RNA-seq data

#### 2.8.1. RNA-seq data processing and gene expression quantification

Raw sequencing reads were subjected to quality control and aligned to the *K. marxianus* FIM1 reference genome (ASM185444v2; GCA_001854445.2) using HISAT2. Gene-level read counts were obtained using featureCounts for subsequent analyses.

#### 2.8.2. Global expression profiling and differential expression analysis

Gene expression data were normalized and variance-stabilized using DESeq2. Principal component analysis (PCA) was performed using the 500 genes with the highest variance in the variance-stabilized expression matrix. Differential expression analysis was performed using DESeq2 with FDHY232 (WT) as the reference. Genes with Benjamini–Hochberg-adjusted *P* values (padj) ≤ 0.05 and |log_2_FC| ≥ 1 were defined as differentially expressed genes (DEGs). All 18 samples were initially evaluated. Because bLYZ_3 showed clear separation from the other bLYZ replicates in the PCA, it was excluded from the primary analysis (n = 17). The complete dataset (n = 18) was retained for sensitivity analysis.

#### 2.8.3. Identification of shared transcriptional responses and candidate genes

UpSet analysis was performed to identify shared and protein-specific DEGs across the five expression groups relative to WT. DEGs were further classified according to their direction of change relative to WT and predefined protein production phenotypes. Candidate genes associated with high- and low-expression phenotypes were identified through six pairwise comparisons between the two phenotype groups. Genes meeting the DEG criteria (padj ≤ 0.05 and |log_2_FC| ≥ 1) in all six comparisons and exhibiting a consistent phenotype-associated direction of change were retained. To identify transcriptional differences between high intracellular expression and high secretion, the rhFTH group was compared with the AnFaeA and BLG-B groups individually. Candidate genes were required to meet the DEG criteria in both comparisons, exhibit consistent directions of change, and show opposite directions of regulation between the intracellular and secretory expression groups relative to WT.

#### 2.8.4. Functional enrichment analysis

Over-representation analysis (ORA) was performed on selected DEG sets using Gene Ontology (GO) annotations, including biological process (BP), cellular component (CC), and molecular function (MF), and KEGG annotations. All detected protein-coding genes were used as the background. Benjamini–Hochberg-adjusted *P* values < 0.05 were considered significant. Gene set enrichment analysis (GSEA) was performed using genes ranked by signed Wald statistics. The n = 17 dataset was used for primary analysis and the n = 18 dataset for sensitivity analysis. Multiple testing correction was performed using the Benjamini–Hochberg method both within and across comparisons. Pathways exhibiting consistent enrichment directions and meeting the false discovery rate (FDR) criteria in both datasets were considered robust.

## 3. Results and discussion

### 3.1. Construction of a K. marxianus chassis combining expression-enhancing genetic traits

To improve chassis compatibility with diverse heterologous proteins, we combined previously identified expression-enhancing genetic traits (Figure 1). T1-Δatg1, which improves heterologous protein production by attenuating autophagy, was selected as the starting strain (19). The *CYR1* N1546K mutation, previously shown to enhance both thermotolerance and heterologous protein production, was subsequently introduced (7). Finally, a translocation between chromosomes III and V, known to enhance protein production by reshaping global cellular metabolism, was introduced using the Cre-loxP system (8) . The resulting chassis strain was designated FDHY232.

**Figure 1.**
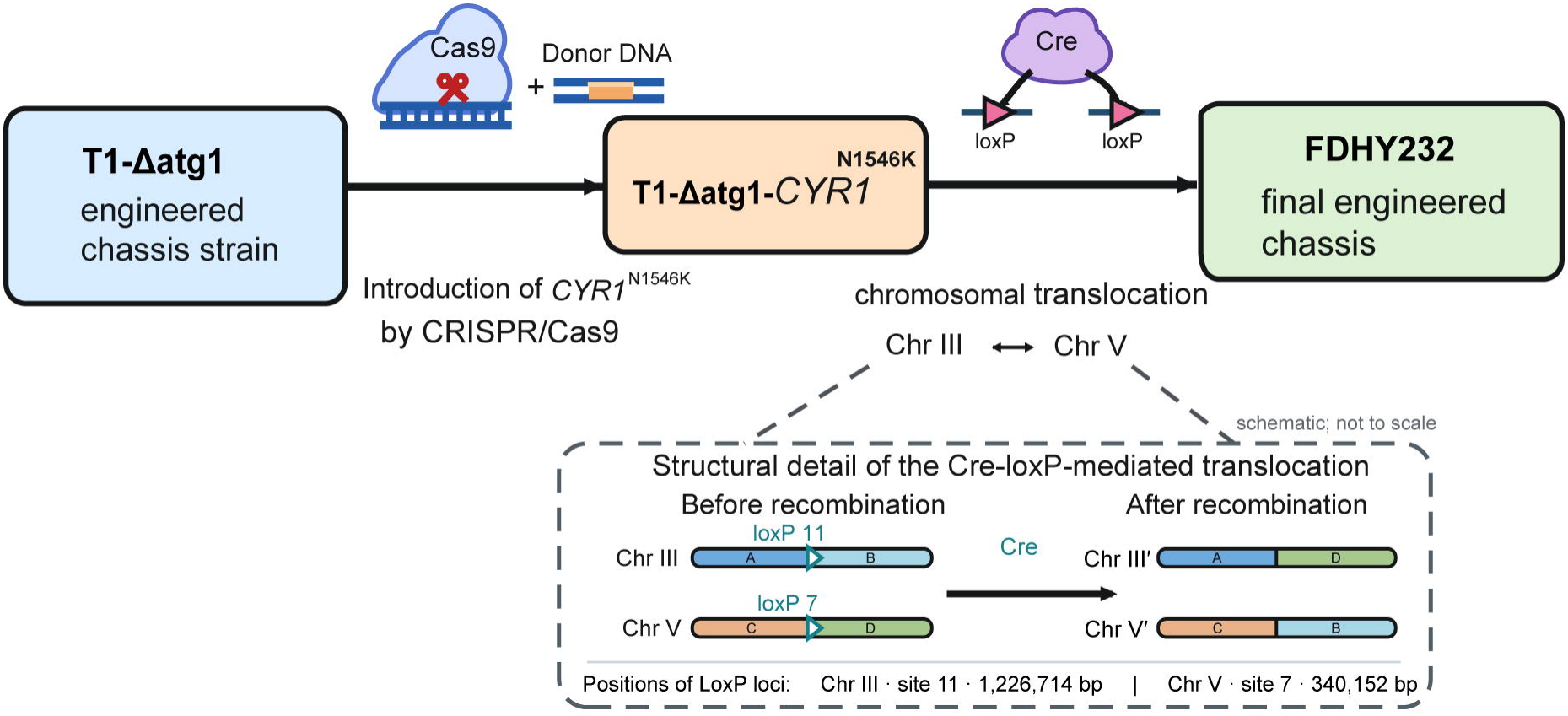
Schematic illustration of the construction of the engineered chassis strain FDHY232. T1-Δatg1 was used as the starting strain. The CYR1 N1546K substitution was introduced by CRISPR/Cas9-mediated genome editing, generating T1-Δatg1-CYR1. Subsequently, Cre-loxP-mediated chromosomal translocation between loxP site 11 on chromosome III (1,226,714 bp) and loxP site 7 on chromosome V (340,152 bp) generated the final engineered chassis FDHY232. The lower schematic illustrates the chromosomal arrangements before and after recombination. Schematic representation is not to scale.

### 3.2. Amino acid composition and characteristics of selected heterologous proteins

To investigate the transcriptional responses of the engineered chassis during the expression of different heterologous proteins, five proteins were selected. Three proteins had been efficiently expressed in KM: human ferritin heavy chain (rhFTH), with an intracellular titer of 11 g/L (20); feruloyl esterase from *Aspergillus niger* (AnFaeA), with a secreted titer of 6 g/L (17) ; and bovine β-lactoglobulin variant B (BLG-B) with substantial secretory expression (21) . Two additional proteins of commercial interest, human lactoferrin (hLF) and bovine lysozyme (bLYZ), which had not yet been efficiently produced in KM, were selected.

Differences in amino acid composition between heterologous and host proteins may affect protein production (27). The relative frequencies of the amino acids in the five heterologous proteins were compared with their distributions across the KM reference proteome (5,070 protein sequences) (Figure 2A). rhFTH and BLG-B exhibited relatively high leucine contents (12.02% and 13.58%, respectively), while AnFaeA contained 10.00% threonine and 8.46% glycine. Among the five proteins, bLYZ had the highest cysteine (6.15%) and tryptophan (4.62%) contents, while hLF contained 4.63% cysteine. The leucine contents of rhFTH and BLG-B, threonine content of AnFaeA, and cysteine contents of bLYZ and hLF exceeded the corresponding medians of the KM reference proteome. These results demonstrate distinct amino acid composition profiles among the five proteins.

**Figure 2.**
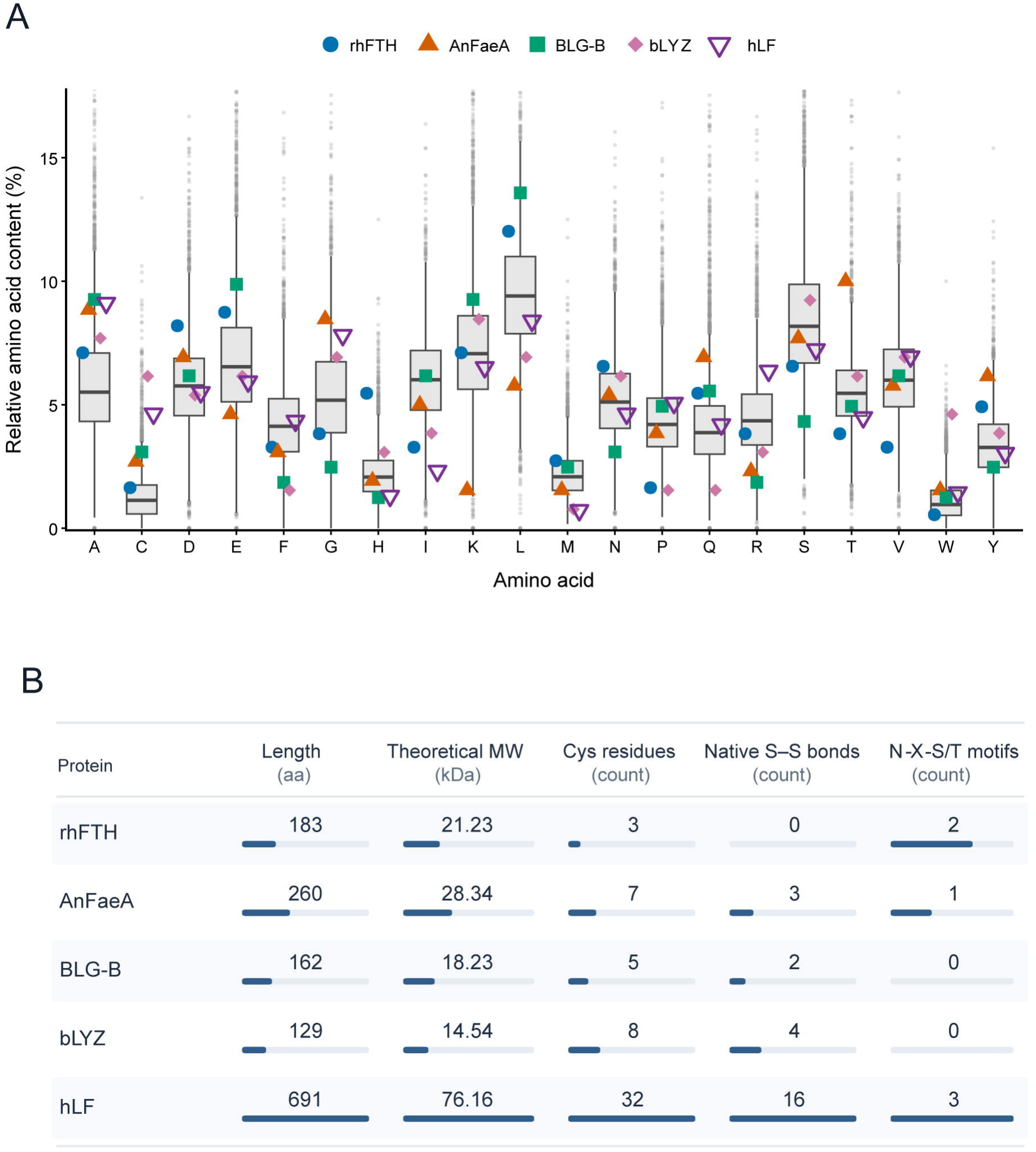
Amino acid composition and molecular characteristics of five heterologous proteins. (A) Comparison of amino acid composition between the five heterologous proteins and the K. marxianus (KM) reference proteome. Boxplots show the distributions of the relative contents of the 20 standard amino acids across 5,070 KM protein sequences. Amino acid contents were calculated as percentages of the total sequence length. Boxes represent interquartile ranges (IQRs), horizontal lines indicate medians, and whiskers extend to the most extreme values within 1.5 × IQR. Gray dots represent outliers, and colored symbols indicate the corresponding values for the five heterologous proteins. The y-axis is restricted to 0–17% for clarity; complete distributions are provided in Supplementary Figure S1. (B) Sequence-derived molecular characteristics and native structural features of the five heterologous proteins, calculated based on their native mature protein sequences. N-X-S/T motif counts represent potential N-linked glycosylation motifs identified from sequence analysis and do not indicate experimentally confirmed glycosylation sites.

Protein size can influence expression efficiency, with smaller proteins generally being more amenable to expression (28). The five proteins varied substantially in theoretical molecular weight (Figure 2B). hLF was the largest (76.16 kDa), whereas bLYZ was the smallest (14.54 kDa). rhFTH, AnFaeA, and BLG-B had theoretical molecular weights of 21.23, 28.34, and 18.23 kDa, respectively.

Disulfide bond number and connectivity are important determinants of protein folding and expression efficiency. hLF contains 32 cysteine residues forming 16 native intramolecular disulfide bonds (29), whereas bLYZ, AnFaeA, and BLG-B contain eight, seven, and five cysteine residues, forming four, three, and two native disulfide bonds, respectively (30-32) (Figure 2B). Consistent with their higher relative cysteine contents, hLF and bLYZ had the greatest numbers of intramolecular disulfide bonds among the five proteins (Fig 2A). In contrast, rhFTH contains three cysteine residues but no known stable intramolecular disulfide bonds in its native structure (33, 34) (Figure 2B). Sequence analysis based on the N-X-S/T motif (X ≠ P) (35) identified three, two, and one potential N-linked glycosylation sites in hLF, rhFTH, and AnFaeA, respectively, but none in BLG-B or bLYZ (Figure 2B).

### 3.3. bLYZ and hLF are poorly expressed in K. marxianus

To evaluate the expression of bLYZ and hLF in FDHY232, three independent transformants expressing each protein (F-bLYZ or F-hLF) were cultivated in shake flasks for 48 h. Culture supernatants and intracellular soluble and insoluble fractions were analyzed by SDS-PAGE and Western blotting, with FDHY232 serving as the parental control (Figure 3).

**Figure 3.**
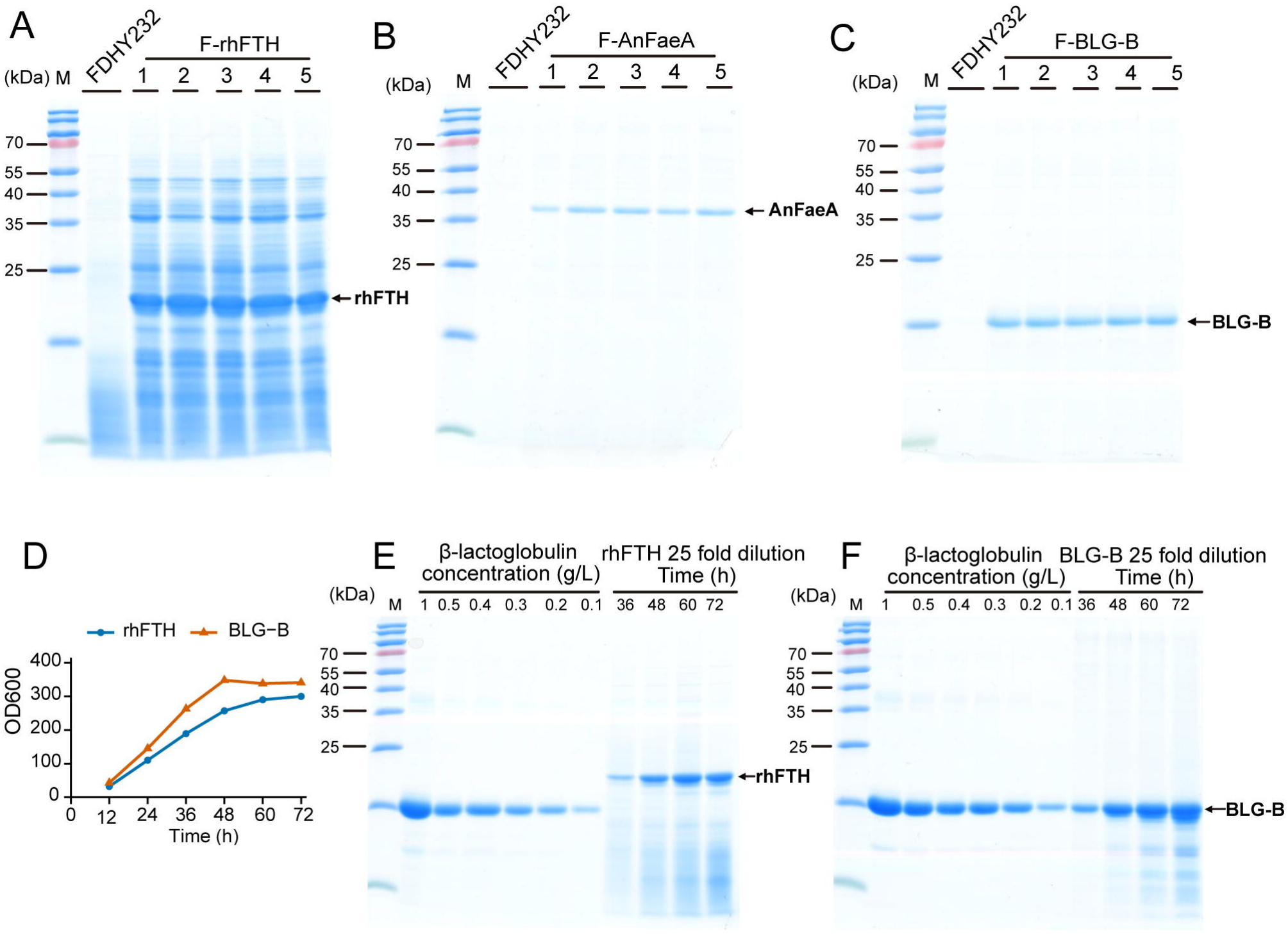
Expression of bLYZ and hLF. (A, B) SDS-PAGE analysis of bLYZ (A) and hLF (B). (C, D) Western blot analysis of bLYZ (C) and hLF (D) using an anti-His antibody. Samples were collected after 48 h of shake-flask cultivation. Extracellular supernatant (E), intracellular soluble fraction (S), and intracellular insoluble fraction (P) were analyzed. Three independent recombinant transformants were examined for each protein, with FDHY232 serving as the parental control.

SDS-PAGE analysis revealed no distinct target protein bands in any of the examined fractions (Figure 3A, B). Western blot revealed distinct bands corresponding to the expected molecular weights of His_6_ tagged bLYZ and hLF in the intracellular insoluble fractions of all three transformants for each protein (Figure 3C, D).

Despite the presence of secretion signal peptides in both expression constructs, bLYZ and hLF predominantly accumulated in intracellular insoluble fractions, suggesting potential limitations in protein folding, processing, or secretion. These results identify bLYZ and hLF as poorly expressed proteins in the engineered KM chassis, providing suitable models for investigating host transcriptional responses associated with inefficient heterologous protein expression.

#### 3.4. rhFTH, AnFaeA, and BLG-B are efficiently expressed in *K. marxianus*

We next evaluated whether rhFTH, AnFaeA, and BLG-B maintained efficient expression levels in FDHY232. After 48 h of shake-flask cultivation, SDS-PAGE analysis revealed prominent rhFTH bands in the intracellular soluble fractions of all five independent transformants (Figure 4A). Similarly, distinct AnFaeA and BLG-B bands were detected in the culture supernatants of all five transformants for each protein (Figure 4B, C), whereas no corresponding bands were observed in the parental control. These results demonstrate efficient intracellular soluble expression of rhFTH and secretion of AnFaeA and BLG-B.

**Figure 4.**
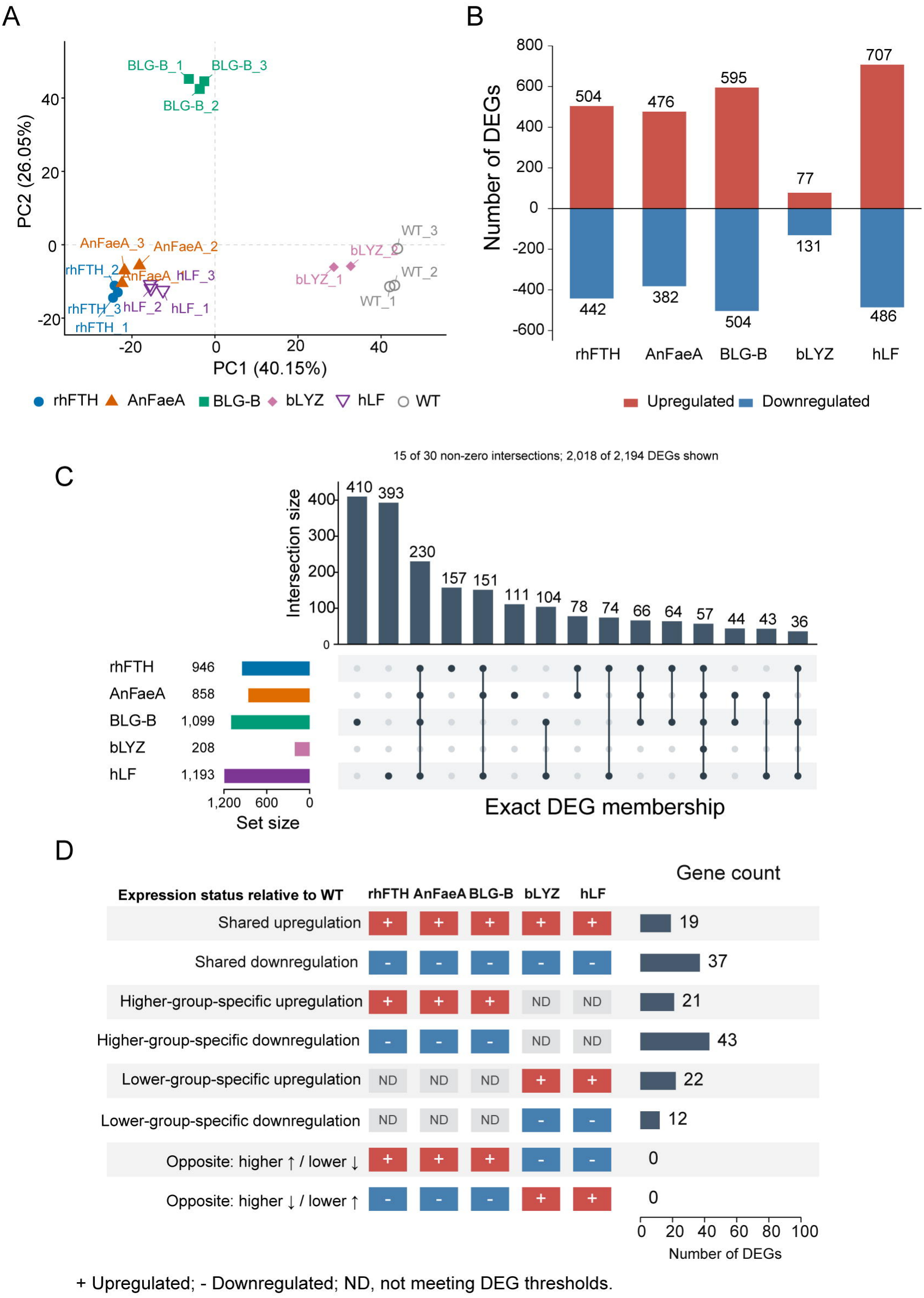
Expression of rhFTH, AnFaeA, and BLG-B. (A–C) SDS-PAGE analysis of intracellular soluble rhFTH (A), secreted AnFaeA (B), and secreted BLG-B (C) after 48 h of shake-flask cultivation. Lanes 1–5 represent five independent transformants, with FDHY232 serving as the parental control. (D) OD_600_ profiles of F-rhFTH and F-BLG-B during fed-batch cultivation. (E, F) SDS-PAGE analysis of intracellular soluble rhFTH (E) and secreted BLG-B (F) during fed-batch cultivation. Protein titers were estimated using β-lactoglobulin standards loaded on the same gels.

To evaluate protein production under high-cell-density conditions, rhFTH- and BLG-B-producing strains were subjected to fed-batch cultivation in 5 L bioreactors. Both strains reached OD_600_ values of approximately 300 or higher at 72 h (Figure 4D). Protein titers were estimated by SDS-PAGE densitometry using β-lactoglobulin as the reference standard. Intracellular soluble rhFTH titers reached 3.62, 10.53, 14.73, and 13.22 g/L at 36, 48, 60, and 72 h, respectively, peaking at 60 h (Figure 4E). Secreted BLG-B titers increased throughout cultivation, reaching 1.47, 5.34, 7.69, and 9.11 g/L at the corresponding time points (Figure 4F).

The maximum estimated rhFTH titer of 14.73 g/L exceeded the previously reported titer of 11.0 g/L in KM (20) and, to our knowledge, represents the highest reported rhFTH titer in a microbial host, highlighting the production potential of FDHY232. Collectively, these results establish rhFTH, AnFaeA, and BLG-B as efficiently expressed proteins suitable for investigating host transcriptional responses associated with efficient heterologous protein expression.

Comparison of the molecular characteristics and expression phenotypes of the five proteins suggests that cysteine content and disulfide bond formation requirements may influence expression efficiency. bLYZ and hLF, which had the highest relative cysteine contents and greatest numbers of native intramolecular disulfide bonds, predominantly accumulated in intracellular insoluble fractions. In contrast, AnFaeA and BLG-B, which had lower cysteine contents and fewer disulfide bonds, were efficiently secreted. Neither molecular weight nor the number of potential N-linked glycosylation sites showed a clear association with expression efficiency (Figure 2).

### 3.5. Global transcriptomic profiles of a K. marxianus chassis expressing different heterologous proteins

To investigate the transcriptional responses of the KM chassis to heterologous protein expression, cells were harvested after 48 h of shake-flask cultivation, with three biological replicates for each of the five protein expression groups and the parental strain FDHY232 (WT). All 18 RNA-seq samples yielded high-quality sequencing data, with Q30 values of 97.33–98.52% and reference genome mapping rates of 81.75–99.17%. Initial PCA revealed that bLYZ_3 deviated substantially from the other two biological replicates (Supplementary Figure S2). Therefore, the dataset excluding bLYZ_3 (n = 17) was used for the primary analysis. PCA of the n17 dataset showed that PC1 and PC2 explained 40.15% and 26.05% of the total variance, respectively (Figure 5A). BLG-B formed a distinct cluster, rhFTH and AnFaeA exhibited similar transcriptional profiles, and bLYZ clustered relatively close to WT. Differential expression analysis of the n17 dataset identified 946, 858, 1,099, 208, and 1,193 differentially expressed genes (DEGs) in the rhFTH, AnFaeA, BLG-B, bLYZ, and hLF groups relative to WT, respectively (Figure 5B; Supplementary Table S1). Notably, hLF and bLYZ, both poorly expressed proteins, exhibited the largest and smallest numbers of DEGs, respectively. These results indicate that different heterologous proteins elicited distinct host transcriptional responses, with no apparent association between the number of DEGs and protein expression efficiency.

**Figure 5.**
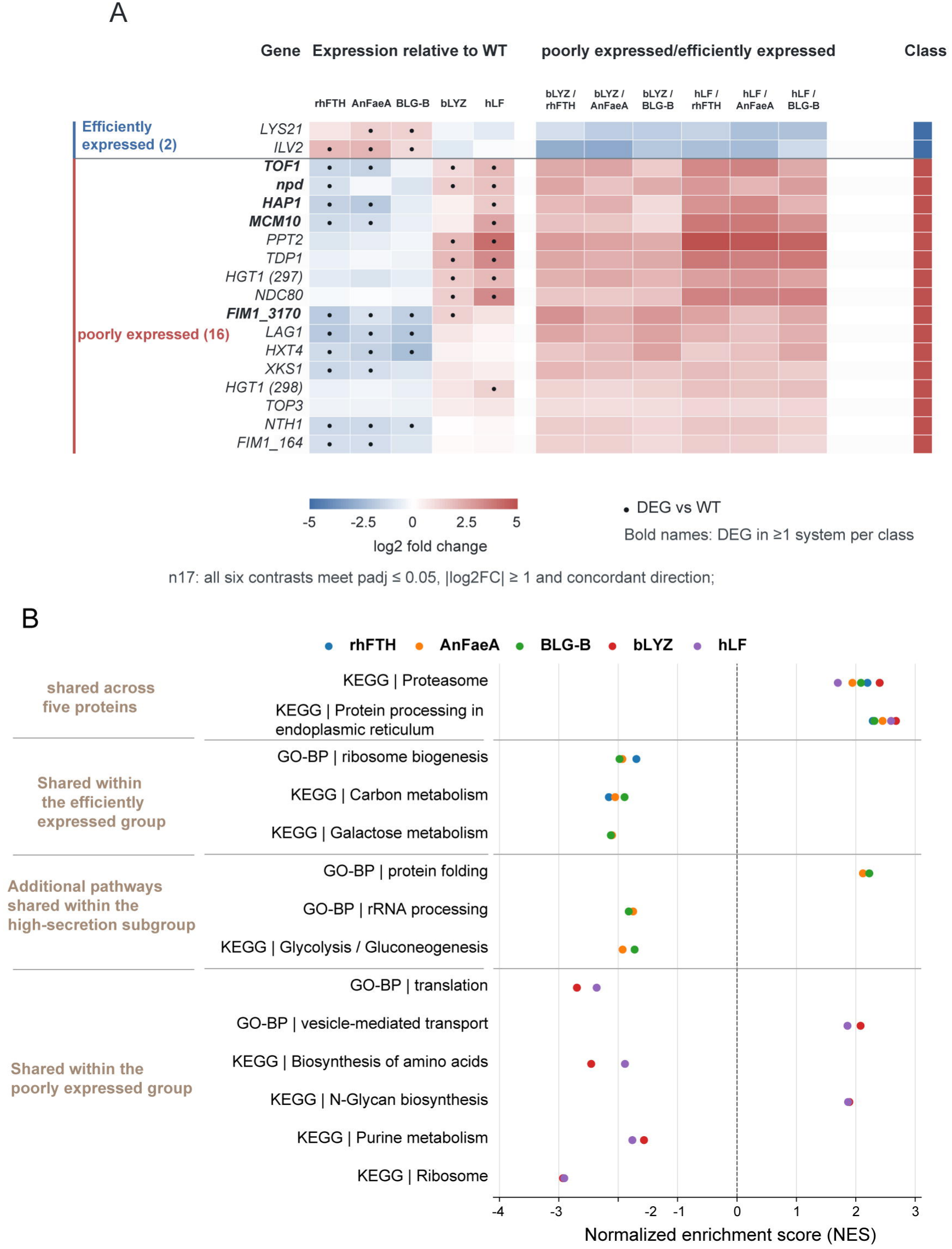
Global transcriptomic profiles of a K. marxianus chassis expressing different heterologous proteins. (A) Principal component analysis (PCA) of 17 RNA-seq samples (n17 dataset) based on the 500 protein-coding genes with the highest variance after variance-stabilizing transformation (VST). PC1 and PC2 explained 40.15% and 26.05% of the total variance, respectively. Colors and shapes distinguish the five protein expression groups and the parental chassis control (WT). (B) Numbers of upregulated and downregulated differentially expressed genes (DEGs) in each expression group relative to WT (padj ≤ 0.05 and |log_2_FC| ≥ 1). (C) UpSet plot showing 15 of the 30 non-empty exact intersections among the five DEG sets, covering 2,018 of the 2,194 non-redundant DEGs. Complete intersections are provided in Supplementary Figure S3. (D) Classification of transcriptional response patterns relative to WT. The efficiently expressed protein group comprises rhFTH, AnFaeA, and BLG-B, whereas the poorly expressed protein group comprises bLYZ and hLF. “+” and “™” indicate upregulation and downregulation, respectively, and “ND” indicates that the DEG criteria were not met. Genes with unavailable statistical results were excluded. This classification reflects transcriptional responses relative to WT rather than direct statistical comparisons between the two phenotype groups.

### 3.6 Shared and distinct transcriptional responses associated with expression phenotypes

Intersection analysis of the five DEG sets identified 2,194 non-redundant DEGs distributed across 30 non-empty exclusive intersections (Figure 5C; Supplementary Figure S3). Among these, 19 genes were consistently upregulated and 37 were consistently downregulated across all five expression systems. These 56 genes accounted for approximately 2.6% of all non-redundant DEGs, indicating a limited shared transcriptional response to heterologous protein expression.

DEGs were further classified according to protein expression phenotypes (Figure 5D). The efficiently expressed protein group (rhFTH, AnFaeA, and BLG-B) exhibited 21 exclusively shared upregulated genes and 43 exclusively shared downregulated genes, whereas the poorly expressed protein group (bLYZ and hLF) exhibited 22 and 12, respectively. No genes met the predefined criteria for opposite regulation between the two phenotype groups. These results suggest that the two groups exhibited distinct shared transcriptional responses rather than simple reciprocal regulation patterns.

Because this classification was based on independent comparisons of each expression system against WT, the identified patterns do not necessarily reflect statistically significant differences between the two phenotype groups. Therefore, direct pairwise comparisons were subsequently performed to identify candidate genes associated with protein expression efficiency.

### 3.7. Candidate genes associated with distinct protein expression phenotypes

To identify transcriptional differences associated with protein expression efficiency, six pairwise comparisons were performed between the poorly expressed protein group (bLYZ and hLF) and the efficiently expressed protein group (rhFTH, AnFaeA, and BLG-B). Eighteen candidate genes met the DEG criteria (padj ≤ 0.05 and |log_2_FC| ≥ 1) and exhibited consistent directions of differential expression across all six comparisons (Figure 6A).

**Figure 6.**
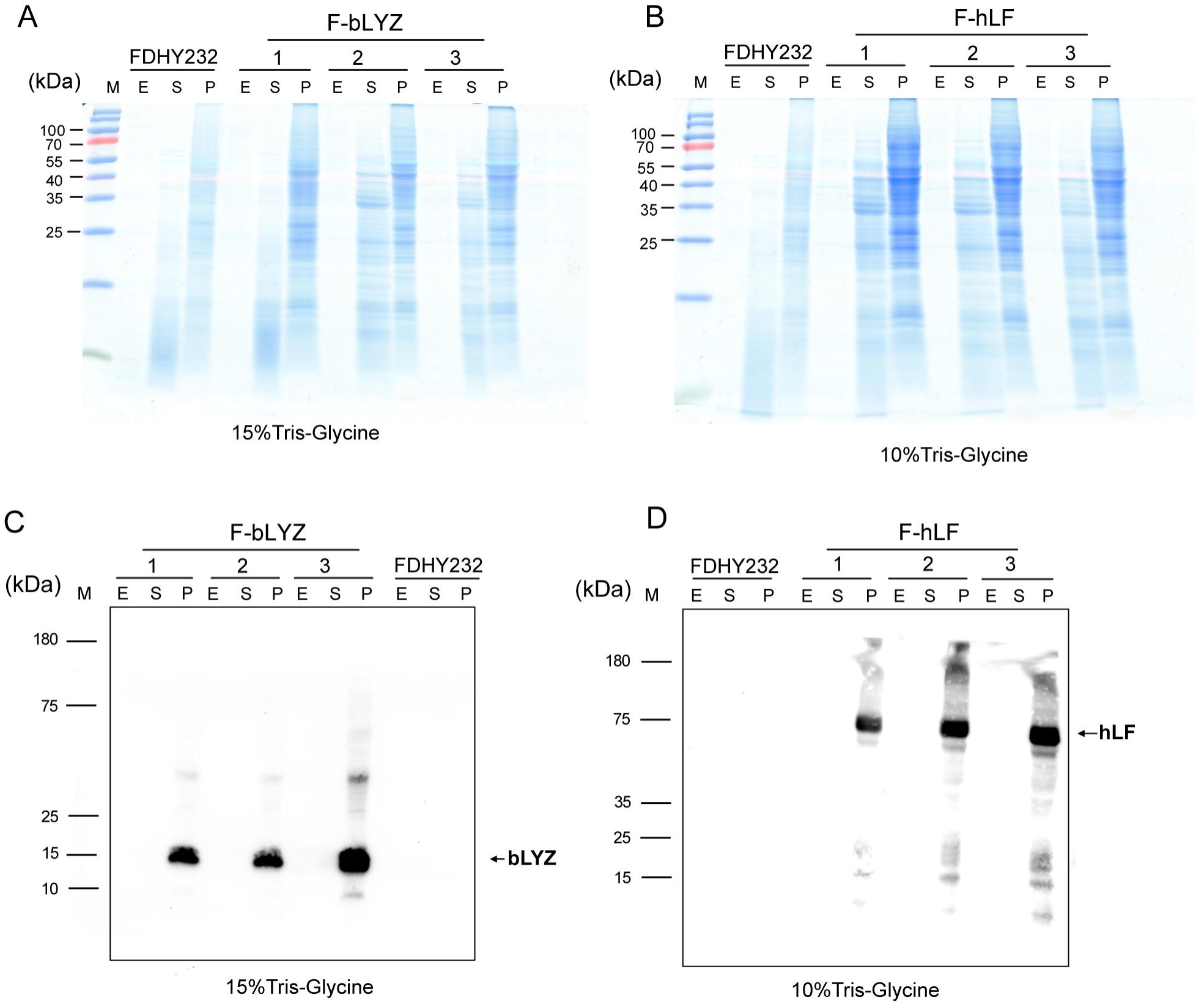
Candidate genes and GSEA pathways associated with distinct heterologous protein-expression phenotypes. (A) Eighteen candidate genes, including two relatively upregulated in the efficiently expressed group and 16 in the poorly expressed group. The left and right heatmaps show log_2_FC relative to WT and six direct pairwise comparisons between the phenotype groups, respectively. Blue and red indicate negative and positive log_2_FC, respectively. Black dots mark DEGs relative to WT, and bold gene names indicate DEGs in at least one system within the corresponding group. Candidates met the predefined criteria (padj ≤ 0.05 and |log_2_FC| ≥ 1) with concordant directions across all six n17 comparisons. HGT1 (297) and HGT1 (298) represent two distinct genomic loci annotated with the same gene name in the FIM1 reference genome and are therefore shown separately. (B) Fourteen representative pathways among the 35 robustly enriched GSEA pathways. Points represent normalized enrichment scores (NES) for the five protein-expression systems. Colors distinguish the systems, and the dashed line indicates NES = 0. Pathways are grouped by shared enrichment patterns.

Of these, *ILV2* and *LYS21* were relatively upregulated in the efficiently expressed protein group, whereas the remaining 16 genes were relatively upregulated in the poorly expressed protein group. *ILV2* and *LYS21* encode acetolactate synthase and homocitrate synthase, which are involved in branched-chain amino acid and lysine biosynthesis, respectively (36, 37). Among the 16 genes, *HAP1* encodes a heme-responsive transcriptional regulator (38), *HXT4* and *XKS1* participate in hexose transport and xylulose metabolism, respectively (39, 40), and *LAG1* is involved in ceramide and sphingolipid biosynthesis (41, 42). These findings suggest that amino acid metabolism, heme-responsive regulation, nutrient transport, and membrane lipid metabolism may be associated with protein expression efficiency, although their causal roles remain to be established.

To distinguish transcriptional responses associated with efficient intracellular expression and secretion, intracellularly expressed rhFTH was individually compared with secreted AnFaeA and BLG-B. A total of 29 candidate features were identified, comprising 28 protein-coding genes and one tRNA-annotated feature. Of these, 12 were relatively upregulated in the rhFTH group and 17 in both secretory expression groups (Supplementary Figure S4). Among these candidates, *LAA1* and *MNN1*, involved in protein trafficking and Golgi glycoprotein modification, respectively, were relatively upregulated in the secretory expression groups (43, 44), consistent with the involvement of these processes in the secretory pathway. In contrast, *EGD1*, involved in co-translational protein folding and targeting (45), and *RPN14*, associated with proteasome assembly (46), were relatively upregulated in the rhFTH group. These findings suggest that nascent polypeptide handling and protein homeostasis may play important roles in efficient intracellular protein production.

### 3.8 Functional enrichment patterns associated with distinct protein expression phenotypes

To identify cellular processes associated with protein expression efficiency, GO and KEGG over-representation analysis (ORA) and gene set enrichment analysis (GSEA) were performed.

ORA revealed significant enrichment of regulation of gene expression among genes consistently regulated across all five expression systems. Commonly upregulated genes in the efficiently expressed protein group were enriched in ER protein processing, whereas commonly downregulated genes were enriched in carbon metabolism, glycolysis/gluconeogenesis, and galactose metabolism. In contrast, commonly downregulated genes in the poorly expressed protein group were significantly enriched in the ribosome pathway (Supplementary Figure S5 and Supplementary Tables S3 and S4).

GSEA identified 35 robust pathways with consistent enrichment directions in the n17 primary and n18 sensitivity analyses, of which 14 representative pathways are shown in Figure 6B. The complete results for all 35 pathways in both datasets are provided in Supplementary Figure S6 and Supplementary Table S2. Protein processing in the ER and the proteasome pathway were positively enriched across all five expression systems, suggesting that remodeling of protein homeostasis represents a common host response to heterologous protein expression, regardless of expression efficiency (Figure 6B and Supplementary Figure S6). This observation is consistent with the coordinated roles of the unfolded protein response (UPR) and ER-associated degradation (ERAD) in maintaining protein homeostasis (47-49). The complete GSEA results for the 35 robust pathways in both datasets are provided in Supplementary Figure S6, and the full statistical results are available in Supplementary Table S2.

Notably, the efficiently and poorly expressed protein groups exhibited distinct transcriptional responses in protein synthesis and metabolism. While negative enrichment of central carbon metabolism and ribosome biogenesis characterized the efficiently expressed protein group, the poorly expressed protein group exhibited negative enrichment of translation, ribosome function, amino acid biosynthesis, and purine metabolism (Figure 6B). These findings suggest that efficient heterologous protein production may require coordinated resource allocation, whereas poor protein expression may be associated with limitations in protein synthesis and precursor availability. Previous studies have demonstrated that enhancing global translation activity can improve recombinant protein secretion in yeast (50). Together with the relatively higher expression of ILV2 and LYS21 in the efficiently expressed protein group (Figure 6A), these results identify translational capacity and amino acid biosynthesis as potential engineering targets for improving poorly expressed proteins.

Another notable difference was the positive enrichment of protein folding in the efficient secretion subgroup, in contrast to the positive enrichment of vesicle-mediated transport and N-glycan biosynthesis in the poorly expressed protein group (Figure 6B). Given the predominant accumulation of bLYZ and hLF in intracellular insoluble fractions and their extensive disulfide bond requirements, the transcriptional responses associated with downstream processing and trafficking may be insufficient to overcome upstream folding limitations. Indeed, engineering disulfide bond formation and ER protein quality control has been shown to enhance heterologous protein production in KM (6). These findings suggest that improving protein folding capacity, particularly oxidative folding and disulfide bond formation, may be more effective than enhancing vesicle-mediated transport alone for these difficult-to-express proteins.

Collectively, the negative enrichment of translation and amino acid biosynthesis pathways in the poorly expressed protein group, together with the differential expression of ILV2 and LYS21, highlights these genes and translation-related genes as potential engineering targets. Moreover, the contrasting enrichment patterns of protein folding and vesicle-mediated transport in the efficient secretion and poorly expressed protein groups suggest that enhancing oxidative folding, rather than further increasing vesicle transport capacity, may improve the production of disulfide-rich proteins.

## Conclusion

Comparative transcriptomic analysis of KM chassis expressing different heterologous proteins identified 18 candidate genes associated with expression efficiency and 29 candidate features distinguishing efficient intracellular expression from secretion. Functional enrichment analysis revealed negative enrichment of translation and amino acid biosynthesis pathways in the poorly expressed protein group and positive enrichment of protein folding in the efficient secretio subgroup. These findings suggest that enhancing amino acid biosynthesis, maintaining translational capacity, and improving oxidative folding are potential engineering strategies for increasing the production of difficult-to-express proteins in KM.

## Acknowledgments

This work was supported by National Key Research and Development Program of China (2021YFA0910601, 2021YFA0910603), and Science and Technology Research Program of Shanghai (24HC2810100, 24ZR1406500, 2023ZX01).

## Data availability

The differential expression summary and complete gene set enrichment analysis (GSEA) results generated in this study are provided in Supplementary Tables S1 and S2, respectively. The authors plan to deposit the raw RNA-seq data in the NCBI Gene Expression Omnibus (GEO). The raw data are not currently publicly available but can be obtained from the corresponding author upon reasonable request.

## Figure legends

**Supplementary Figure S1**. Full-range comparison of amino acid composition between five heterologous proteins and the Kluyveromyces marxianus reference proteome. Relative contents of the 20 standard amino acids are shown across the full 0–100% range. Boxplots represent the distributions across 5,070 KM proteins, with center lines indicating medians, boxes indicating interquartile ranges (IQRs), and whiskers extending to 1.5 × IQR. Colored symbols indicate the corresponding values for the five heterologous proteins.

**Supplementary Figure S2**. Principal component analysis of the complete RNA-seq dataset (n = 18). PCA was performed using the 500 protein-coding genes with the highest variance after variance-stabilizing transformation (VST). Each point represents an individual biological sample, with colors and shapes distinguishing the five protein-expression groups and the parental chassis control (WT). Sample labels indicate individual replicates. PC1 and PC2 explained 44.55% and 21.41% of the total variance, respectively.

**Supplementary Figure S3**. Complete DEG intersections among five heterologous protein-expression systems in the n17 primary analysis. UpSet plot showing all 30 non-empty exact intersections among the five DEG sets identified by comparison with WT. The horizontal bars indicate the total number of DEGs in each expression system, while the vertical bars represent the sizes of individual intersections. Connected dots indicate the exact combinations of DEG sets represented by each intersection. DEGs were defined by padj ≤ 0.05 and |log_2_FC| ≥ 1.

**Supplementary Figure S4**. Candidate features associated with efficient intracellular expression versus high secretion in the n17 primary analysis. The heatmap shows the expression changes of 29 candidate features relative to WT in the rhFTH, AnFaeA, and BLG-B systems, together with two direct pairwise comparisons (rhFTH versus AnFaeA and rhFTH versus BLG-B). Twelve features were relatively upregulated in the rhFTH group, whereas 17 were relatively upregulated in both secretory expression groups. Black dots indicate DEGs relative to WT. The n18 column indicates whether each candidate retained the predefined selection criteria in the complete dataset. Filled and open circles indicate retained and non-retained candidates, respectively. The 29 features comprise 28 protein-coding genes and one tRNA-annotated feature.

**Supplementary Figure S5**. GO and KEGG over-representation analysis of phenotype-associated gene sets in the n17 primary analysis. Enriched terms are shown for genes consistently regulated across all five expression systems, commonly upregulated and downregulated genes in the efficiently expressed protein group, and commonly downregulated genes in the poorly expressed protein group. Dot size represents gene count, dot color indicates ™log_10_ of the Benjamini–Hochberg-adjusted P value, and the x-axis represents the gene ratio. Only selected significantly enriched GO terms and KEGG pathways (adjusted P < 0.05) are shown.

**Supplementary Figure S6**. Complete GSEA results for 35 robustly enriched pathways in the n17 primary and n18 sensitivity analyses. Panel A shows normalized enrichment scores (NES) from the n17 primary analysis (17 samples), and panel B shows the corresponding results from the complete n18 dataset (18 samples). Each point represents the NES for a pathway in a protein-expression system, with colors distinguishing the five systems. The dashed vertical line indicates NES = 0. Pathways are grouped according to shared enrichment patterns across all five systems, the efficiently expressed group, the high-secretion subgroup, and the poorly expressed group. Robust pathways exhibited consistent enrichment directions and met the FDR criteria in both datasets.

