## Supplementary figures and images for "Comparative Transcriptomic Analysis of a Kluyveromyces marxianus Chassis Expressing Heterologous Proteins with Different Expression Efficiencies"

### Supplementary Figure S1

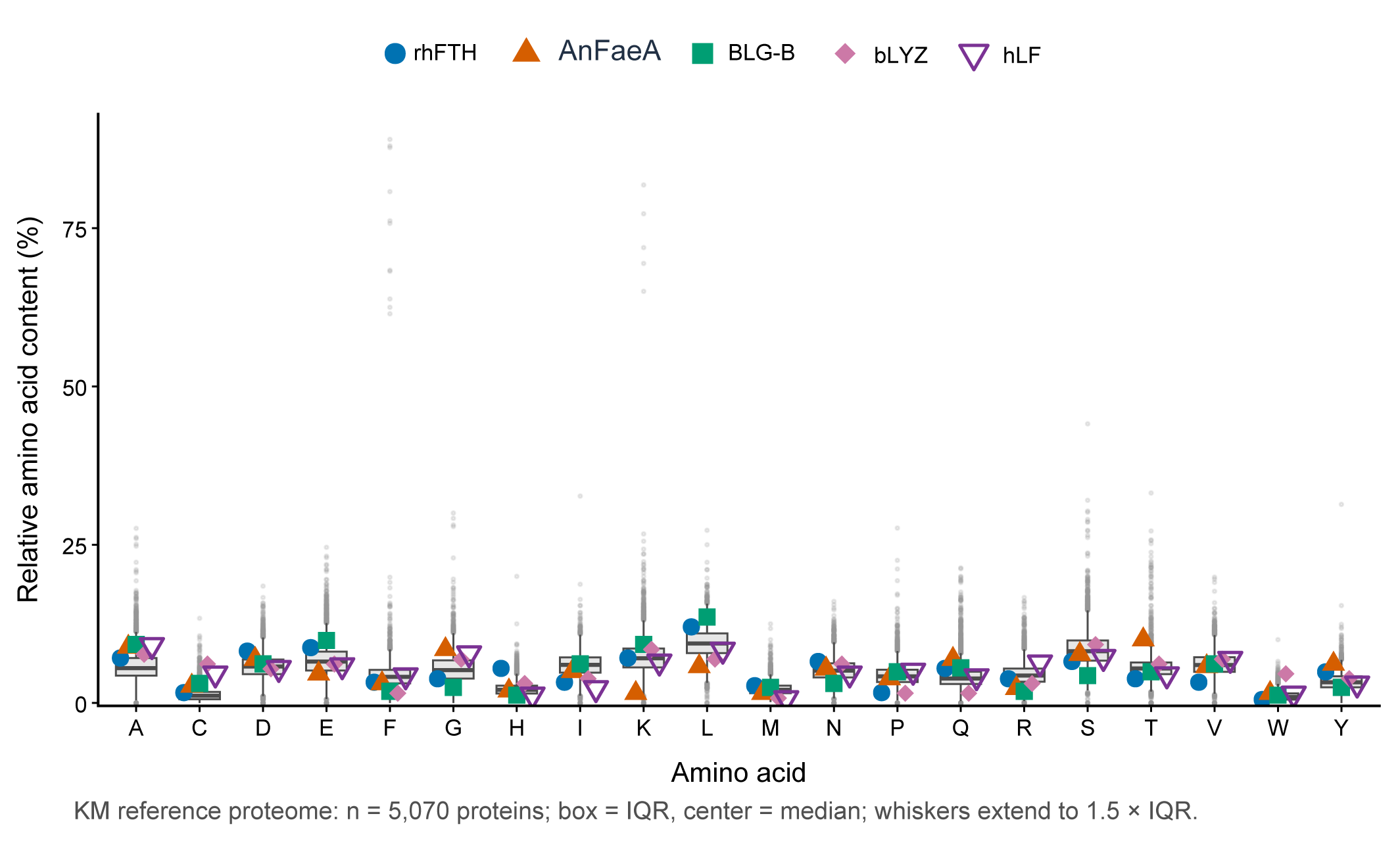

### Supplementary Figure S2

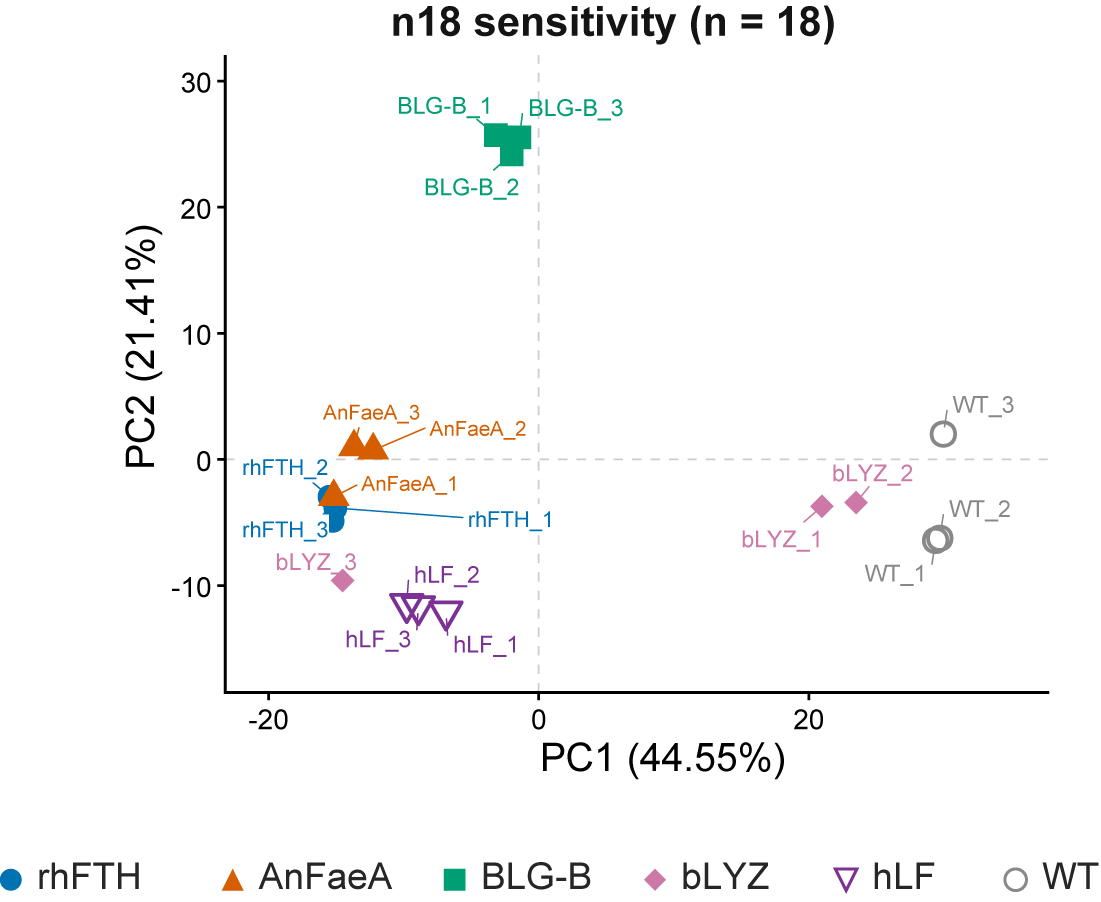

### Supplementary Figure S3

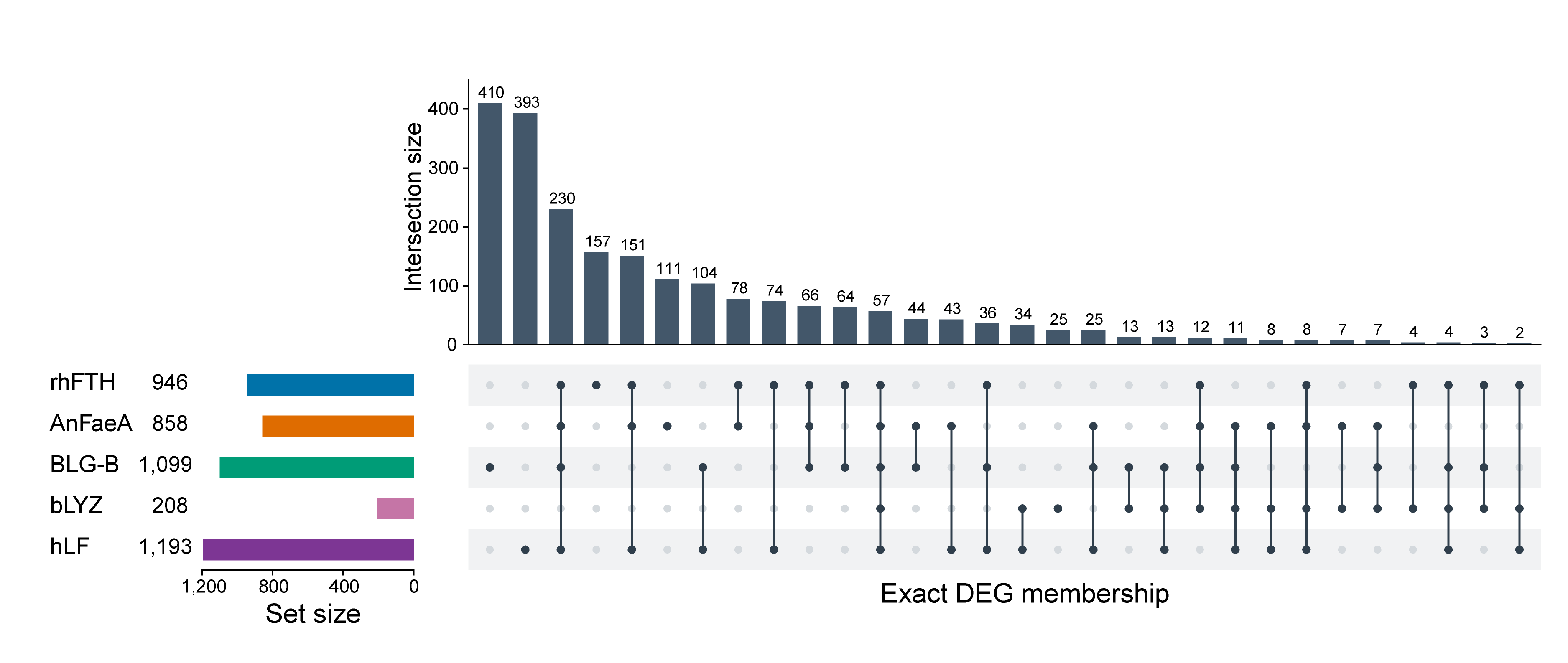

### Supplementary Figure S4

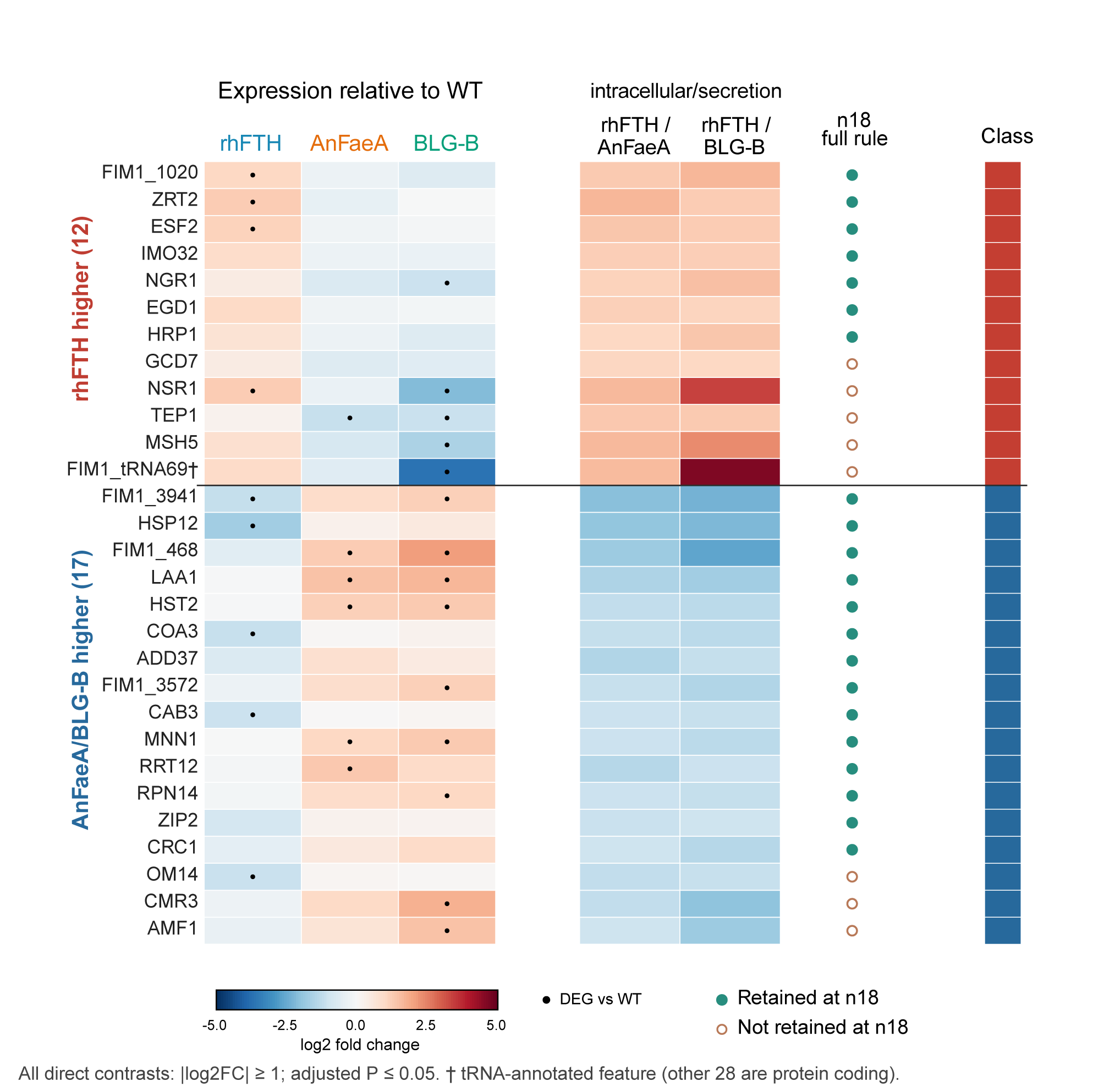

### Supplementary Figure S5

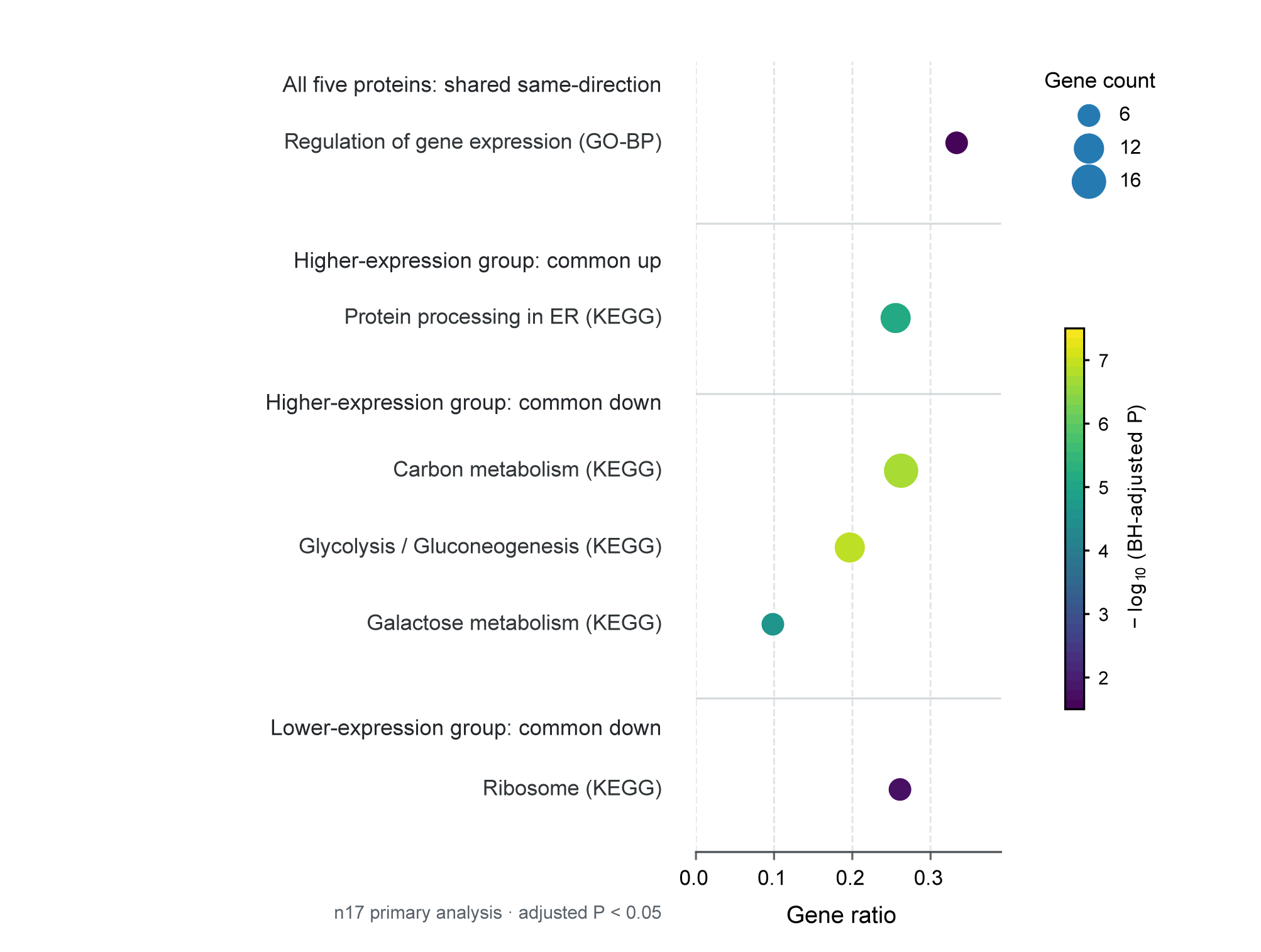

### Supplementary Figure S6

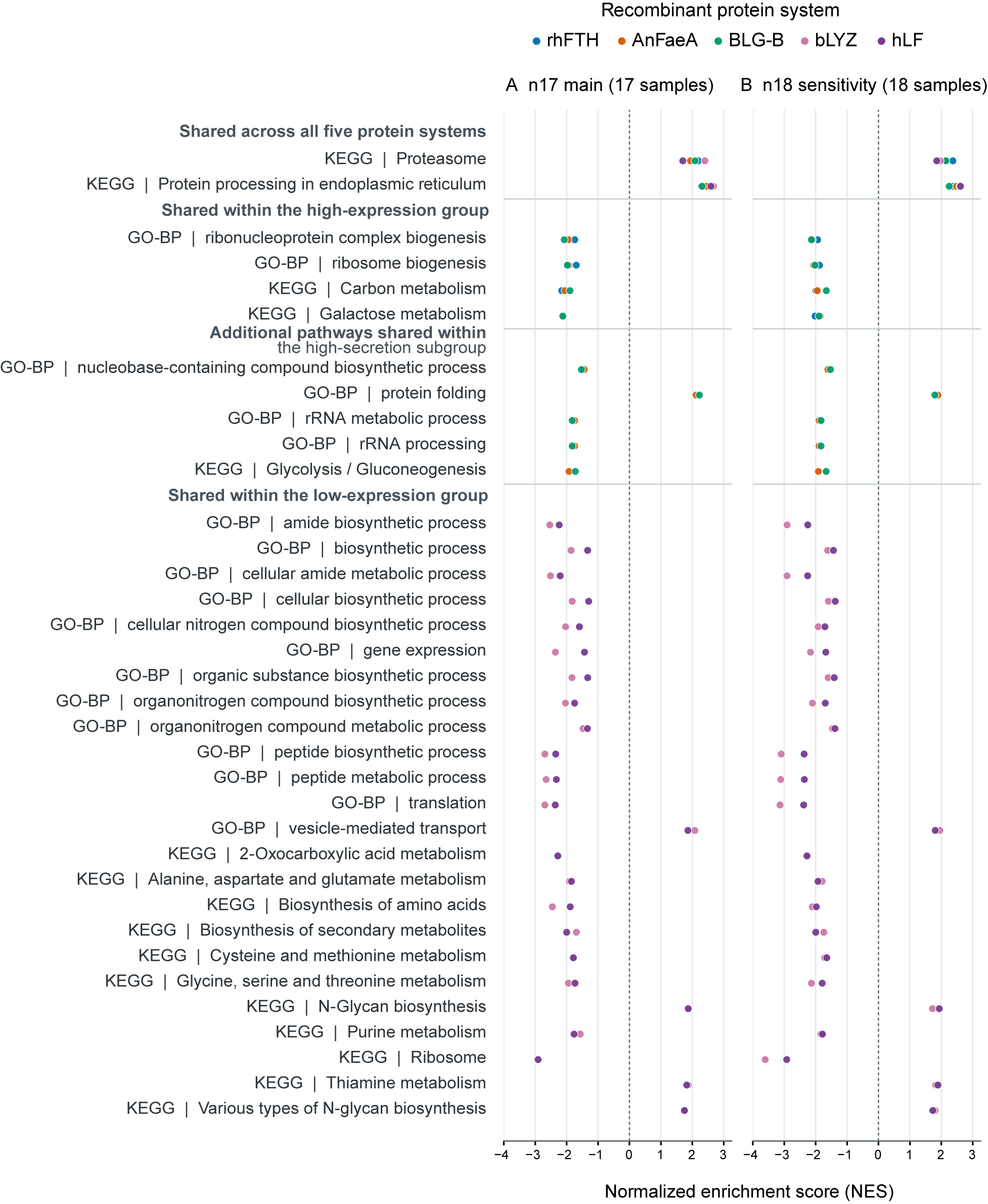
