## Supplementary File 1 for "Comparative Transcriptomic Analysis of a Kluyveromyces marxianus Chassis Expressing Heterologous Proteins with Different Expression Efficiencies"

The codon-optimized sequences of bLYZ and hLF.

| Protein Name | Nucleic acid sequence |
| --- | --- |
| Bovine lysozyme（bLYZ） | AAGGTTTTCGAAAGATGTGAATTGGCTAGAACTTTGAAGAAGTTGGGTTTGGATGGTTACAAGGGTGTTTCTTTGGCTAATTGGTTGTGTTTGACTAAGTGGGAATCTTCTTACAATACTAAGGCTACTAATTACAATCCATCTTCTGAATCTACTGATTACGGTATTTTCCAAATTAATTCTAAGTGGTGGTGTAATGATGGTAAGACTCCAAATGCTGTTGATGGTTGTCATGTTTCTTGTAGAGAATTGATGGAAAATGATATTGCTAAGGCTGTTGCTTGTGCTAAGCATATTGTTTCTGAACAAGGTATTACTGCTTGGGTTGCTTGGAAGTCTCATTGTAGAGATCATGATGTTTCTTCTTACGTTGAAGGTTGTACTTTGCATCACCATCACCATCAT |
| Human lactoferrin（hLF） | GGTAGAAGGAGATCTGTTCAATGGTGTGCAGTTTCTCAGCCTGAAGCTACTAAGTGTTTCCAATGGCAAAGAAATATGAGAAAGGTTAGAGGTCCACCTGTTTCTTGTATTAAGAGAGATTCTCCAATTCAATGTATTCAAGCTATTGCTGAAAATAGAGCTGATGCTGTTACTTTGGACGGTGGTTTTATTTACGAAGCTGGTTTGGCTCCATACAAGTTGAGACCTGTTGCTGCTGAAGTTTACGGTACTGAAAGACAACCAAGAACTCATTACTACGCTGTTGCTGTTGTTAAGAAAGGTGGTTCTTTCCAATTGAATGAATTGCAAGGTTTGAAGTCTTGTCATACTGGATTAAGAAGAACTGCTGGTTGGAACGTTCCAATTGGTACTTTGAGACCATTCTTGAATTGGACTGGTCCACCTGAACCAATTGAAGCTGCTGTTGCTAGATTCTTCTCTGCTTCTTGTGTTCCTGGTGCTGATAAGGGTCAATTCCCAAATTTGTGTAGATTGTGTGCTGGTACTGGTGAAAACAAATGTGCTTTCTCTTCTCAAGAACCATACTTCTCTTACTCTGGTGCATTCAAGTGTTTGAGAGATGGTGCTGGAGATGTTGCTTTTATTAGAGAATCTACTGTTTTCGAAGATTTGTCTGATGAAGCTGAAAGAGATGAATACGAATTGTTGTGTCCTGATAATACTAGAAAGCCTGTTGATAAGTTCAAGGATTGTCATTTGGCTAGAGTTCCATCTCATGCTGTGGTTGCTAGATCTGTTAATGGTAAGGAAGATGCTATTTGGAATTTGTTGAGACAAGCTCAAGAAAAGTTCGGTAAGGATAAGTCTCCAAAGTTCCAATTGTTCGGTTCTCCATCTGGTCAAAAGGATTTGTTGTTCAAGGATTCTGCTATTGGTTTCTCTAGAGTTCCACCAAGAATTGATTCTGGTTTGTACTTGGGTTCTGGTTACTTCACTGCTATTCAAAATT |

Continued.

| Protein Name | Nucleic acid sequence |
| --- | --- |
|  | TGAGAAAGTCTGAAGAGGAAGTTGCTGCTAGAAGAGCTAGAGTTGTTTGGTGTGCTGTTGGTGAACAAGAATTGAGAAAGTGTAATCAATGGTCTGGTTTGTCTGAAGGTTCTGTTACTTGTTCTTCTGCTTCTACTACTGAAGATTGTATTGCTTTGGTTTTGAAGGGTGAAGCTGATGCTATGTCATTGGACGGTGGTTATGTGTACACAGCTGGTAAGTGTGGTTTGGTTCCTGTTTTGGCTGAAAACTATAAATCTCAACAATCTTCTGATCCTGATCCAAATTGTGTTGATAGACCTGTTGAAGGTTACTTGGCTGTTGCTGTTGTTAGAAGGTCTGATACTTCTTTGACTTGGAATTCTGTTAAGGGTAAGAAGTCATGTCATACAGCTGTTGATAGAACAGCTGGATGGAATATTCCTATGGGTTTATTATTCAATCAAACTGGTTCTTGTAAGTTCGATGAATACTTCTCTCAATCTTGTGCTCCTGGTTCTGATCCAAGATCTAATTTGTGTGCTTTGTGTATTGGTGATGAGCAAGGTGAAAATAAGTGTGTTCCAAATTCTAATGAAAGATACTACGGTTACACTGGAGCTTTCAGATGTTTGGCTGAGAATGCTGGTGATGTTGCTTTCGTTAAGGATGTTACTGTTTTGCAAAATACTGATGGTAATAACAATGAAGCTTGGGCTAAGGATTTGAAGTTGGCTGATTTCGCTTTGTTGTGTTTGGATGGTAAGAGAAAGCCTGTTACAGAAGCTAGATCTTGCCATTTGGCTATGGCTCCAAATCACGCTGTTGTTTCTAGAATGGATAAGGTTGAAAGATTGAAGCAAGTTTTGTTGCATCAACAAGCTAAGTTCGGTAGAAATGGTTCTGATTGTCCTGATAAGTTCTGTTTGTTCCAATCTGAAACTAAGAATTTATTGTTCAACGACAACACTGAATGTTTGGCTAGATTGCATGGTAAGACTACTTACGAAAAGTACTTGGGTCCACAATACGTTGCTGGTATTACTAATTTGAAGAAGTGTTCTACTTCTCCATTGTTGGAAGCTTGTGAATTCTTGAGAAAGCATCACCATCACCATCAT |
